# Genome and transcriptome of a pathogenic yeast, *Candida nivariensis*

**DOI:** 10.1101/2021.01.27.428461

**Authors:** Yunfan Fan, Andrew N Gale, Anna Bailey, Kali Barnes, Kiersten Colotti, Michal Mass, Luke B Morina, Bailey Robertson, Remy Schwab, Niki Tselepidakis, Winston Timp

## Abstract

We present a highly contiguous genome and transcriptome of the pathogenic yeast, *Candida nivariensis*. We sequenced both the DNA and RNA of this species using both the Oxford Nanopore Technologies (ONT) and Illumina platforms. We assembled the genome into an 11.8 Mb draft composed of 16 contigs with an N50 of 886 Kb, including a circular mitochondrial sequence of 28 Kb. Using direct RNA nanopore sequencing and Illumina cDNA sequencing, we constructed an annotation of our new assembly, supplemented by lifting over genes from *Saccharomyces cerevisiae* and *Candida glabrata*.

## Introduction

For immunocompromised hosts, opportunistic infections caused by drug resistant fungi of the *Candida* genus are a major source of morbidity and mortality(Borman *et al*. 2008). In particular, *Candida nivariensis*, a close relative to *Candida glabrata*, has emerged in recent years as especially resistant to antifungal therapies(Borman *et al*. 2008). However, due to its phenotypic similarities to *C. glabrata, C. nivariensis* is generally underidentified and easily misdiagnosed, and currently, only molecular approaches can distinguish the two (Aznar-Marin *et al.* 2016), spurring whole genome sequencing studies on the clade(Gabaldón *et al.*. 2013)

Accurate assembly of repetitive genomic regions is crucial for understanding genetic diversity and virulence in pathogenic species. Fungal pathogens have long been known to exhibit a high degree of genome plasticity to enhance fitness in various environments(Croll *et al.* 2013; Ford *et* al. 2015; López-Fuentes *et al.* 2018; Carreté *et al.* 2019; Todd *et al.* 2019). Repetitive subtelomeric regions in particular play a crucial role in virulence for many pathogenic organisms(Barry *et al.* 2003; De Las Peñas *et al.* 2003). Many yeasts’ subtelomeric regions contain and regulate the expression of genes crucial for biofilm formation, carbohydrate utilization, and cellular adhesion(Naumov *et al.* 1995; De Las Peñas *et al.* 2003; Iraqui *et al*. 2005). These gene families often undergo rapid evolution through changes in copy number and sequence through either SNPs or indels(Carreto *et al.* 2008; Brown *et al.* 2010; Anderson *et al*. 2015). However, these subtelomeric regions remain one of the most difficult sections of the genome to accurately assemble due to their repetitive nature and high sequence similarity between genes, making genetic analysis cumbersome(Brown *et al.* 2010).

One of the gene families that are of great interest to the pathogenic yeast field is the GPI-anchored cell wall proteins. This protein family includes many genes that encode for adhesion proteins that are found in various members of the Candida genus and play a key role in pathogenicity; such as regulate biofilm formation, cell-to-cell contact, and host-pathogen interactions(Timmermans *et al.* 2018; McCall *et al.* 2019). With the many roles these genes play in infection the accurate identification and understanding of the genetic variation of these genes vital to combating fungal pathogens.

Unfortunately, like many eukaryotic pathogens, the current reference genome for C. *nivariensis* is highly fragmented, in 123 contigs with an N50 of 248Kb, meaning that at least half of the total genome length is contained in contigs 248Kb or longer. This is typical of genomes assembled from limited short-read sequencing data; though short reads are highly accurate, assembling them into contiguous genomes is challenging depending on the size and complexity of the genome. Such short read assemblies have limited utility since large scale variants, repetitive regions, and genome structure remain difficult to elucidate, though they are often involved in the genome plasticity of pathogenic yeasts(Carreté *et al.* 2018). In contrast, long read sequencing data has been shown to produce much more contiguous assemblies, and have been crucial in sequencing through large repetitive regions, as well as assessing structural variants. However, read accuracy on the ONT platform in particular ranges from 86% for early basecaller versions(Wick *et al.* 2019) to 97% as currently reported by ONT. This is lower than the read accuracy of short read Illumina sequencing, which achieves 99.9% accuracy(Fox *et al.* 2014). In consensus sequences, most random errors can be corrected by other reads covering the same genomic loci, resulting in >99% consensus accuracy(Wick *et al.* 2019). However, systematic errors occurring in most or all of the reads cannot be corrected this way. For ONT data, indels at homopolymers dominate systematic errors(Wick *et al.* 2019). These persistent errors can be problematic for gene prediction and annotation in downstream analysis(Watson and Warr 2019), and are typically corrected with more accurate short read data in mappable regions (Garrison and Marth 2012; Walker *et al.* 2014; Vaser *et al.* 2017)

Having a genome alone is not enough; we need to annotate it with genes and other functional elements for the genome to be of greatest use. Knowledge of gene loci is critical to constructing phylogenetic relationships between organisms, and to studying the functional implications of variants, both common uses of reference genomes. While model-based, purely computational gene predictors can be highly accurate in bacteria, gene sparsity and intronic regions make this task more difficult in eukaryotes(Salzberg 2019). For improved annotations, some RNA-seq information is required(Salzberg 2019).

Here, as part of our newly developed Methods in Nucleic Acid Sequencing class, we used a hybrid approach, applying long read nanopore sequencing to assemble a highly contiguous genome of *C. nivariensis*, followed by short read sequencing to polish or correct errors in our assembly. We followed this by a combination of nanopore direct RNA sequencing as well as short read RNA-seq to annotate our assembly. By combining this data with liftover of annotations from evolutionary “cousins” of nivariensis, we have generated a new and annotated reference genome for the community.

## Materials and Methods

### Media and growth conditions

For genomic extractions, a single colony of *C. nivariensis* CBS9983, originally isolated from a blood culture of a Spanish woman(Alcoba-Flórez *et al.* 2005), was inoculated into synthetic complete (SC) medium supplemented with 2% glucose and shaken overnight at 30°C in a glass culture tube. For RNA extractions, *C. nivariensis* CBS9983 was grown to log phase in SC medium supplemented with 2% glucose at 30°C in a glass culture tube.

### DNA isolation and sequencing

DNA was extracted from liquid culture using the Zymo Fungal/Bacterial DNA MiniPrep Kit according to manufacturer specifications. Two ONT sequencing libraries were prepared from the extracted DNA using the ONT rapid barcoding sequencing kit (SQK-RBK004), and each was sequenced on a separate MinION flowcell (R9.4). Two Illumina libraries were prepared with the Nextera Flex Library Prep Kit, each using 400 ng of extracted DNA. Both Illumina libraries were then sequenced on a single iSeq 100 run.

### RNA Sequencing

RNA was extracted from liquid culture using the Zymo Fungal/Bacterial RNA MiniPrep Kit. Using the NEBNext Poly(A) mRNA Magnetic Isolation Module, polyA tailed mRNA was isolated from the total RNA. Two ONT direct RNA sequencing libraries were prepared and sequenced on separate MinION flowcells, each using ∼200 ng of polyA selected RNA and the SQK-RNA002 sequencing kit. With the NEBNext Ultra II RNA First Strand Synthesis Module and the NEBNext Ultra II Non-Directional RNA Second Strand Synthesis Module, cDNA was prepared from the isolated mRNA. Two individual Illumina libraries were then prepared with the Nextera Flex Library Prep Kit, each using 400 ng of cDNA. Both library replicates were then sequenced on a single iSeq 100 run.

## Results

### Genome assembly

Nanopore data was basecalled using Guppy v3.2.4 on default settings. Reads greater than 3kb long with an average basecalling quality score greater than 7 were assembled into 21 contigs using Canu v2.1 (Koren *et al.* 2017) on default settings. Illumina DNA reads were trimmed for adapters and quality using Trimmomatic v0.39(Bolger *et al.* 2014) using settings `LEADING:3 TRAILING:3 SLIDINGWINDOW:4:30 MINLEN:36`. The trimmed reads were then used to iteratively correct draft assembly using Freebayes(Garrison and Marth 2012) with alignments made by bwa mem(Li 2013) using default settings. Changes were made at positions with alternative allele frequency greater than .5 and the total number of alternate allele observations was greater than 5. We aligned and corrected the assembly iteratively for three rounds, after which further rounds of corrections made no changes.

Of our 21 corrected contigs, 5 were flagged as repeats by Canu and originally constructed from fewer than 180 nanopore reads. The remaining 16 contigs were constructed from over 1800 nanopore reads each. Because the 5 repetitive contigs were constructed from so comparatively few reads, we excluded them from the final assembly. One 32 Kb contig was suggested to be circular by Canu, and therefore likely to be a mitochondrial sequence. To confirm, we aligned this contig to the complete mitochondrial genome of *Candida nivariensis* (NCBI sequence NC_036379.1) using Mummer(Marçais *et al.* 2018), and observed a 3662bp sequence in the reference mitochondrial genome which appears at both ends of our 32kb circular contig. Using the Mummer alignments (Supp. Figure 1), we removed the extraneous 3662bp from the end of our contig, resulting in a 28 kb mitochondrial genome, which we named ‘JHU_Cniv_v1_mito.’ Lastly, we remapped the ONT and Illumina reads back to the assembly, and found no bases with zero coverage, indicating that none of our contigs need to be further broken (Supp. Figure 2). The resulting final assembly comprises 11.8 Mb of sequence in 16 contigs with an N50 of 886 Kb (Figure 1a, Table 1).

**Table 1:** Assembly statistics.

|  | Contigs | N50 | Longest Contig | Shortest Contig | Total Length |
| --- | --- | --- | --- | --- | --- |
| Reference | 123 | 248 Kb | 807 Kb | 666 bp | 11.56 Mb |
| JHU_Cniv_v1 | 16 | 886 Kb | 1.42 Mb | 28.5 Kb | 11.83 Mb |

**Figure 1:**
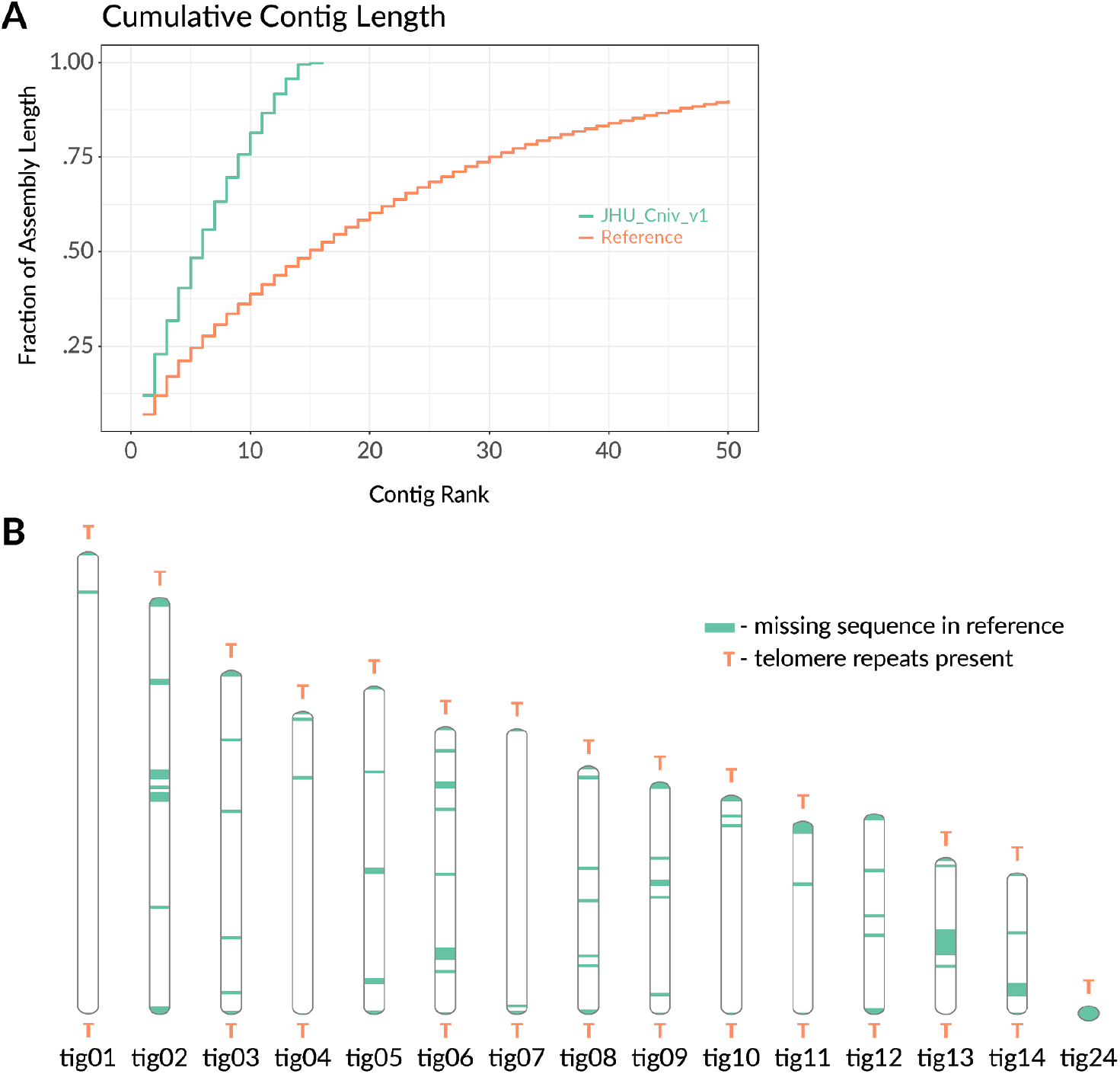
Characteristics of the JHU_Cniv_v1. (a) Cumulative lengths of the 50 longest sequences in our assembly and previous reference genome. (b) Ideogram of assembly. Sequence that is missing in the reference genome is shown along each non-mitochondrial contig, and the positions of telomere repeats are marked.

Of the 275kb of additional sequence in our assembly compared to the current reference, 218kb are accounted for by gaps in the reference which are newly spanned by our JHU_Cniv_v1 Of the 69 newly spanned gap sequences, 54 were identified as repeat regions by Tandem Repeats Finder(Benson 1999) (TRF) with settings [match = 2, mismatch = 7, delta = 7, pm = 80, pi = 10, minscore = 50, maxperiod = 600](Xu *et al.* 2020). Another 17 gap regions were identified to contain a high proportion (10%) of multi-mapping short reads as identified by bwa mem(Li 2013) on default settings.

To determine whether JHU_Cniv_v1 contigs represent full chromosomes, we looked for telomere repeats in our assembly and attempted to use related yeast reference genomes to scaffold. In our assembly, 11 contigs terminate at both ends in repeats of CTGGGTGCTGTGGGGT, the telomere sequence of *Candida glabrata(McEachern and Blackburn 1994)*. The other 4 non-mitochondrial sequences terminate only at one end in this telomeric repeat (Figure 1b, Supp. Table 1), suggesting they may scaffold to form two additional chromosomes. This suggests that, like *C. glabrata*, the *C. nivariensis* genome also contains 13 chromosomes.

We tried to further scaffold our assembly using the more contiguous and highly related *glabrata* genome as a reference, but we found that reference based scaffolders such as Medusa(Bosi *et* al. 2015) and RagTag(Alonge *et al.* 2019) either placed telomeric sequences in the middle of scaffolds or made no improvement (Supp. Figure 3). Upon aligning the *C. glabrata* genome to JHU_Cniv_v1 using Mummer, we found only sporadic shared segments of negligible length (Supp. Figure 4), as opposed to a nearly perfect 1:1 alignment between JHU_Cniv_v1 and the current C. *nivariensis* reference genome (Supp. Figure 5). This indicates that the C. *glabrata* genome is not sufficiently similar to C. *nivariensis* to use as a reference for contig scaffolding. Using the C. *nivariensis* reference genome itself for scaffolding results in similar mis-scaffolds or no change to our assembly, which is unsurprising, as the C. *nivariensis* reference genome is so highly fragmented.

To assess assembly completeness, fungal single copy orthologues were checked using BUSCO (Simão *et al.* 2015) and its available saccharomycetes_odb10 database. Out of 2137 BUSCOs searched, JHU_Cniv_v1 has only 14 missing, 13 of which are also missing in the current reference (Figure 2). This missing gene, RNA polymerase archaeal subunit P/eukaryotic subunit RPABC4 (buscoID 41996at4891), though present in the reference, has the second lowest combined match length and match score among all genes searched. From the reference, we extracted the nucleotide sequence of this match using the coordinates reported by BUSCO, and searched for it in JHU_Cniv_v1 using blastn. We found a full length match with 99.9% identity, suggesting that this BUSCO is not actually absent in JHU_Cniv_v1. Upon further examination of this alignment, we found that all 7 non-matching nucleotides consist of small deletions associated with poly-A or poly-T homopolymers, known error-prone regions for nanopore sequencing data(Watson and Warr 2019).

**Figure 2:**
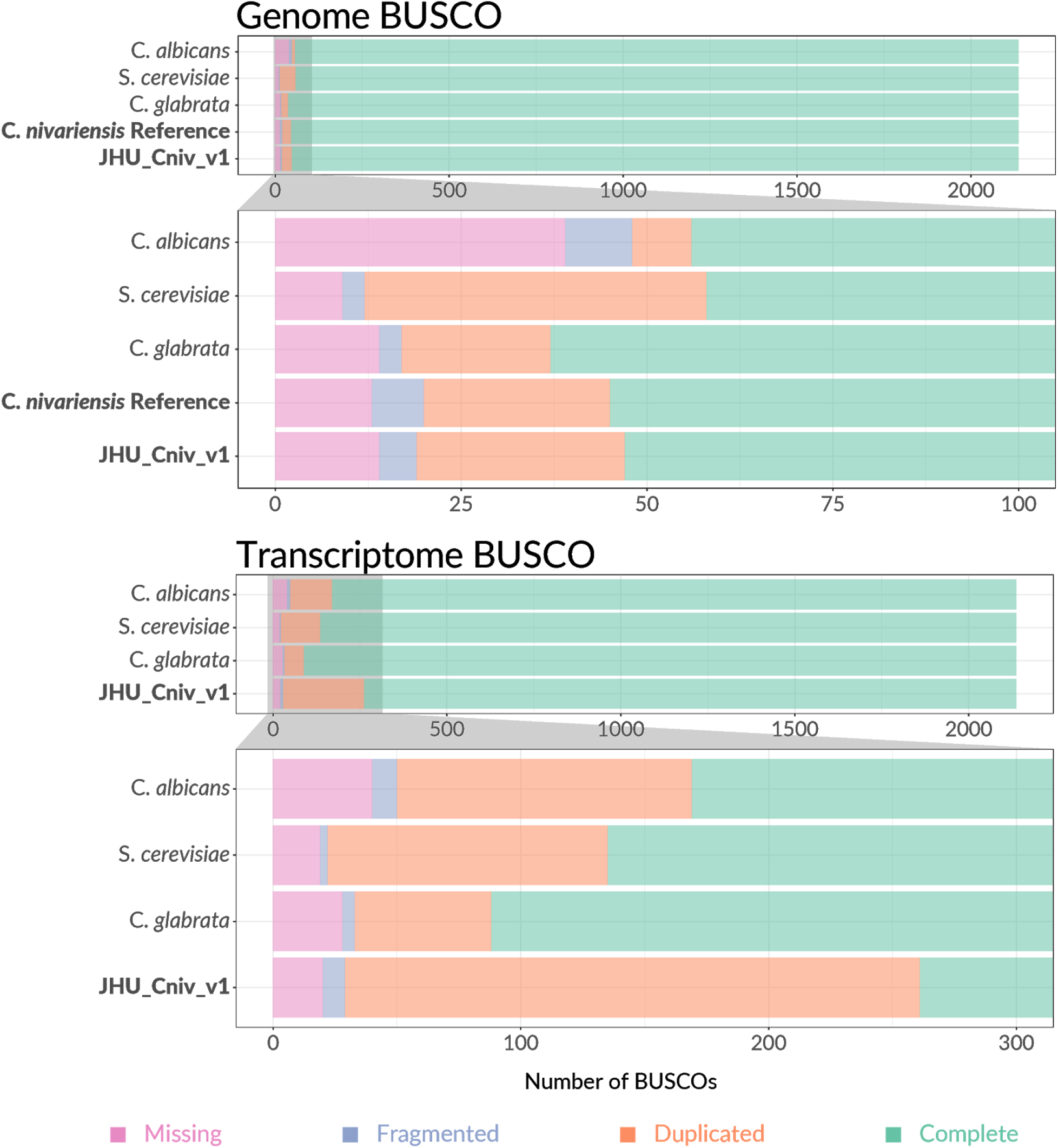
Genome and transcriptome completeness. Bar charts comparing BUSCOs detected in JHU_Cniv_v1 and accompanying transcriptome to those of the current C. albicans, S. cerevisiae, C. glabrata, and C. nivariensis reference genomes. No reference annotation is currently available for C. nivariensis.

### Annotation

Illumina RNA-seq reads were trimmed using Trimmomatic v0.39(Bolger *et al.* 2014) in order to check for any remaining adapter sequences, and to filter out reads with low base quality. HISAT2 was used on default settings to align the trimmed cDNA reads to the assembly. The BRAKER(Hoff *et al.* 2019) pipeline was then used to make gene predictions using these alignments. Currently, ONT dRNA compatibility with BRAKER is in development, and that data was thus not used for prediction. Instead, ONT dRNA reads were aligned to the genome assembly using Minimap2 v2.17(Li 2018) on recommended settings for nanopore direct RNA reads (`-ax splice -uf -k14`). Transcripts were then assembled from the dRNA alignments using Stringtie2(Kovaka *et al.* 2019) with the long read option (`-L`). Using Liftoff(Shumate and Salzberg 2020), we lifted over the annotations from *C. glabrata, S. cerevisiae, C. albicans*.

Starting with the BRAKER predictions, Gffcompare(Pertea and Pertea 2020) was used to add non-overlapping annotations lifted from *C. glabrata, S. cerevisiae*, and *C. albicans* in that order. Specifically, we add any annotation with class code “u” in the Gffcompare .tmap outputs when comparing our list of genes with a list of potential genes to add. Finally, we compared and added non-redundant transcripts assembled by stringtie2 to the annotation using gffcompare. Our final annotation comprises 25,979 features, 5,859 of which are genes (Supp. Table 2). Current annotations of closely related yeasts report similar gene counts (Supp. Table 3).

In order to assess transcriptome completeness, BUSCO was used in transcriptome mode, again with the saccharomycetes_odb10 database. Because no annotation of the *C. nivariensis* reference genome currently exists, we compared our transcriptome to those of *C. glabrata, S. cerevisiae*, and *C. albicans*. Compared to these highly characterized yeast transcriptomes, ours contains slightly fewer complete and single copy BUSCOs (1876 of 2137 searched) and roughly double the number of complete and duplicated BUSCOs (232 of 2137 searched). The numbers of missing and fragmented BUSCOs between the three are comparable (Figure 2).

### Repetitive genes

As *C. glabrata* subtelomeric regions have been proven to be difficult to correctly assemble using short read data(Xu *et al.* 2020), we compare the copy number of *C. glabrata* subtelomere gene homologs between the *C. nivariensis* reference genome and JHU_Cniv_v1. Using the assembly and re-annotation of *C. glabrata* and the from Xu et al.(Xu *et al.* 2020), we extracted the sequences of the C. *glabrata* subtelomere genes and used BLAST (blastn v2.6.0+) to find any matches in the C. *nivariensis* reference and JHU_Cniv_v1. We observed an identical set of 48 C. *glabrata* subtelomere genes in both C. *nivariensis* genomes, but found that the copy number for several were greater in JHU_Cniv_v1 (Figure 3A). To account for genes truncated by short contigs in the reference genome, we calculate copy number by summing the alignment lengths of all the hits of a particular gene and dividing by gene length. Of the 48 *C. glabrata* genes with homology in C. *nivariensis*, 35 are ribosomal. With the exception of just three ribosomal genes which occur a similar number of times in both C. *nivariensis* genomes, all homologous ribosomal genes appear once in the reference, and either four or six times in JHU_Cniv_v1 (Figure 3A, Supp. Data 1).

**Figure 3:**
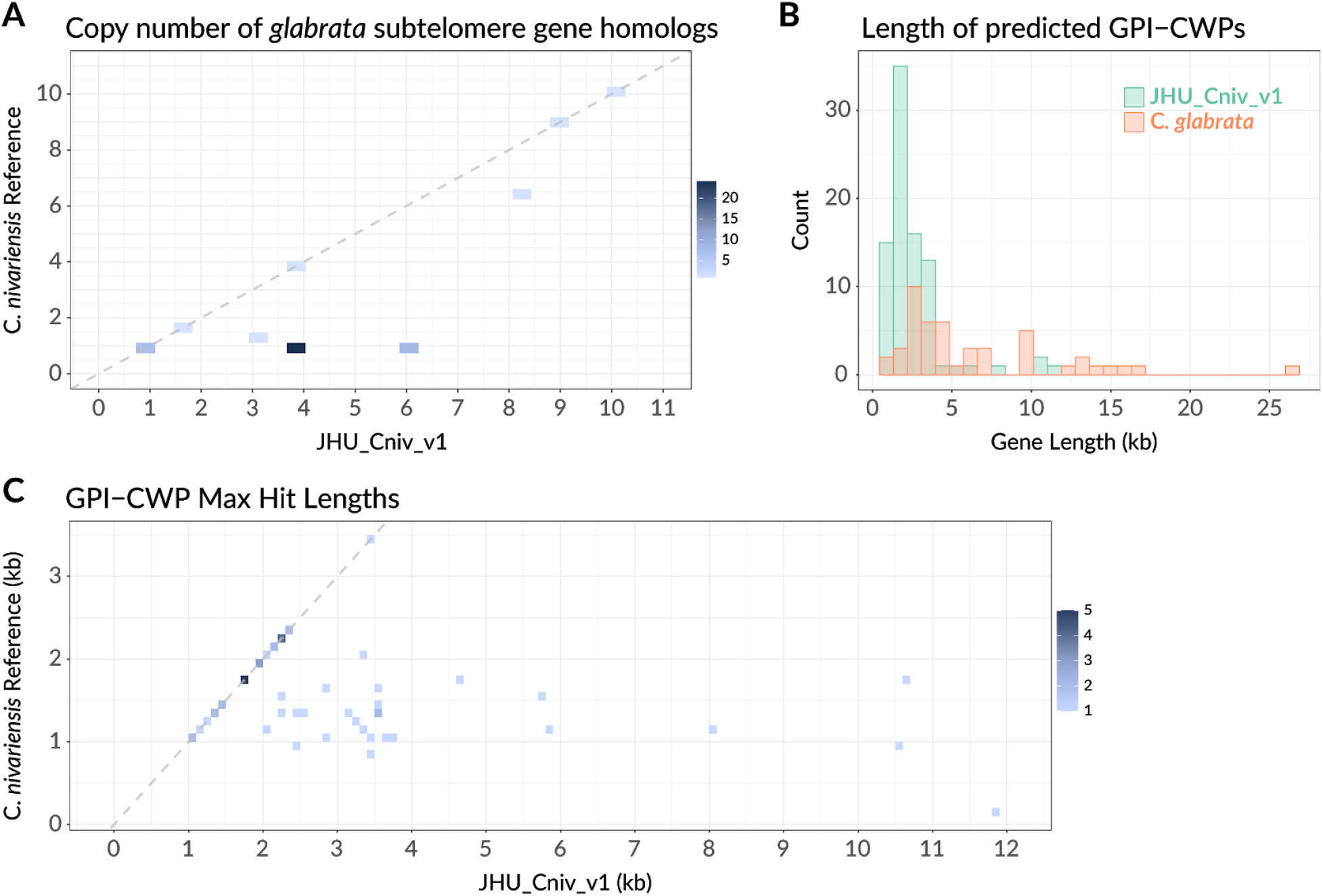
GPI genes. (a) Copy number of glabrata subtelomere gene homologues in the C. nivariensis reference genome and JHU_Cniv_v1. The y=x line is shown in dashed grey. (b) Histogram of adhesion protein lengths in glabrata as annotated by Xu et al, and the lengths of predicted adhesion proteins found in JHU_Cniv_v1. (c) Maximum BLAST alignment lengths for each predicted nivariensis GPI gene in JHU_Cniv_v1 and the C. nivariensis reference genome. The y=x line is shown in dashed grey.

Using JHU_Cniv_v1, we identified GPI-anchored membrane proteins among annotated genes >1000 nt long. Using GffRead(Pertea and Pertea 2020), we constructed the amino acid sequences for these genes and excluded any with internal stop codons. We then used PredGPI(Pierleoni *et al.* 2008) to predict which of these encoded GPI proteins, using an FDR cutoff of <.0005(Xu *et al.* 2020) to find 86 total genes. As GPI anchored fungal adhesins typically contain tandem repeats(Lipke 2018; Xu *et al.* 2020), we further filtered for genes overlapping with tandem repeats as classified by TRF, and identified 53 of the GPI genes as putative adhesins. As with *C. glabrata*, the putative adhesins typically spanned multiple kilobases (Figure 3b), though we do not find very long (>13kb) genes in contrast to several *glabrata* GPI-CWPs. To find the corresponding adhesin genes in the *C. nivariensis* reference genome, we again used blastn, and compared the longest hit of adhesin gene to the true length of the gene as predicted in JHU_Cniv_v1 (Figure 3c). Notably, no hit in the reference genome exceeded 3.5kb, and 27 of these adhesin genes are not found continuously, suggesting the previous reference either truncated or did not continuously assemble these important pathogenic genes.

## Discussion

JHU_Cniv_v1 is a high quality, extremely contiguous assembly of *Candida nivariensis* constructed by long reads and polished by short reads. It spans large, repetitive gaps in the *nivariensis* genome that have fragmented short read assemblies thus far, and includes a full mitochondrial chromosome, as well as telomere repeats. These telomere repeats are identical to those in C. *glabrata*, and have been found to be shared within the entire ‘glabrata group(Gabaldón *et al.* 2013).’ The orientation of the telomeres suggests that C. *nivariensis* has 13 chromosomes, which is in agreement with previous PFGE data(Gabaldón *et al.* 2013). Furthermore, of the contigs missing telomere repeats on one end, we note that scaffolding tig05 with tig12 and tig02 with tig24 would result in 13 chromosomes that would all match PFGE length estimates to 8% error or less, which is within the expected range of PFGE error for very large DNA fragments(Cutting *et al.* 1988).

As assessed by BUSCO, genome completeness of the current C. *nivariensis* reference and JHU_Cniv_v1 is comparable to other related yeasts and is slightly improved over the previous reference. However, while JHU_Cniv_v1 is a much more contiguous assembly than any *C. nivariensis* genome preceding it, the few remaining sequence errors still can pose a problem to downstream analyses, as evidenced by the seemingly absent BUSCO we manually identified.

Our accompanying RNA-seq data enabled us to annotate this genome, achieving a similar level of BUSCO completeness to some of the most highly studied model organisms. Our annotation has comparable or lower levels of missing and fragmented BUSCOs compared to the reference annotations, though more duplicated ones. While our annotation is largely comparable to those of similar yeasts, it has not been manually curated, and should thus be treated as preliminary. Of course, as these organisms were grown under only one condition before RNA extraction, it remains unlikely that this annotation is fully complete.

To demonstrate the utility of genome and annotation contiguity, we examine genes from a difficult to assemble region in C. *glabrata*. For each subtelomeric C. *glabrata* gene with homology in C. *nivariensis*, more copies were found in JHU_Cniv_v1, as its contiguity allows it to more easily capture repeated genome elements. We note that of subtelomeric glabrata genes found, the majority are ribosomal, and of these, only three do not show a four or six times increased copy number in JHU_Cniv_v1. Due to the repetitive nature of rDNA arrays, it can be difficult for short read genome assemblies to capture them in their full complexity. Conversely, our long read assembly more easily spans these regions, potentially providing a clearer look at the biology in which they are involved.

In addition to genes arranged in complex and repetitive patterns, our more contiguous assembly enables analysis of large genes with internal repeats, such as GPI adhesins. Since these genes are so large, it can be difficult or impossible to predict them from fragmented assemblies which are unable to capture them in their full length. As adhesins are critical to understanding elements of pathogenicity in these yeasts, fragmented genome assemblies and missing gene annotations can be crippling to this dimension of research in these organisms.

## Acknowledgements

Much of the data described in this paper was collected as part of the EN.580.454 Methods in Nucleic Acid Sequencing course at Johns Hopkins University in Spring of 2019. We thank Illumina Inc. and ONT for supporting educational initiatives and supplying reagents for collecting this data.

This work was funded in part by the Johns Hopkins University Department of Biomedical Engineering, National Institute of Health (NIH) grant 1R01HG009190 (WT), and Canadian Institutes of Health Research (CIHR) Doctoral Foreign Study Award DFS-157831 (YF). WT holds two patents (US 8,748,091 and US 8,394,584) licensed to ONT. YF and WT have received travel funding from ONT.

## Data Availability

All sequence data is available in the Sequence Read Archive (SRA), under BioProject PRJNA686979. The JHU_Cniv_v1 assembly is available at accession JAEVGP000000000. Code used for analysis is available at https://github.com/timplab/nivar.

