## Supplemental Material for "Genome and transcriptome of a pathogenic yeast, *Candida nivariensis*"

**SUPPLEMENTARY MATERIAL**

### TABLE OF CONTENTS

**Supplemental Figures:**

**Supplemental Figure 1:** Whole genome alignment of the JHU_Cniv_v1 mitochondrial contig and the C. *nivariensis* mitochondrial genome.

**Supplemental Figure 2:** Coverage histograms

**Supplemental Figure 3:** Telomere positions reference based scaffolds

**Supplemental Figure 4:** Whole genome alignments between related yeasts

**Supplemental Figure 5:** Whole genome alignment of JHU_Cniv_v1 and the C. *nivariensis* reference genome

**Supplemental Tables:**

**Supplemental Table 1:** Contig lengths and contig telomere counts

**Supplemental Table 2:** Contributions from each annotation software

**Supplemental Table 3:** Gene and exon counts of JHU_Cniv_v1 and related yeasts

**Supplemental Data:**

**Supplemental Data 1:** Copy numbers of subtelomeric homologues

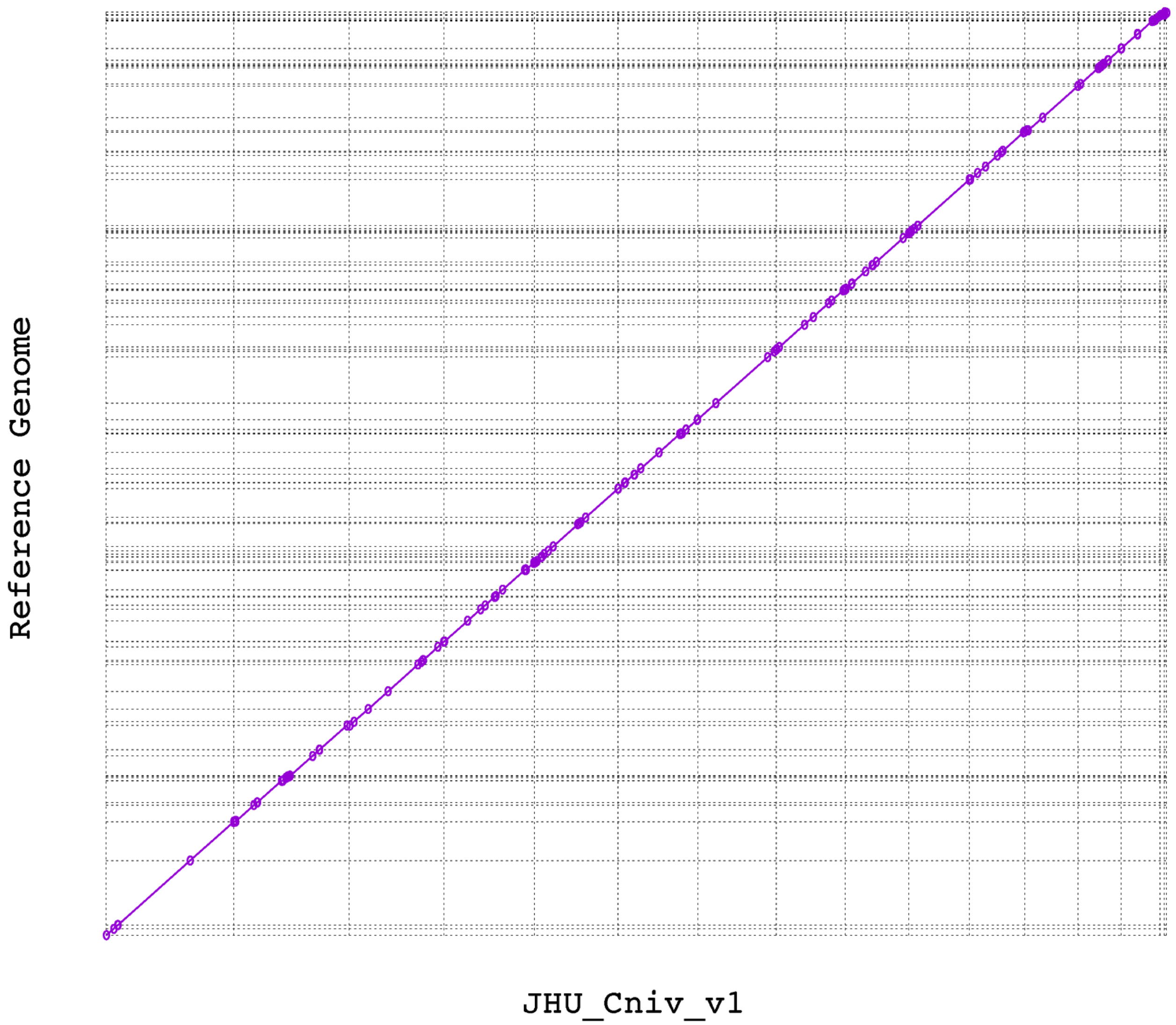


**Supplemental Figure 1**: Whole genome alignment of the current reference genome (y axis) compared and our new assembly (x axis). Alignments match with no notable structural variants, and very little missing or duplicated sequence.

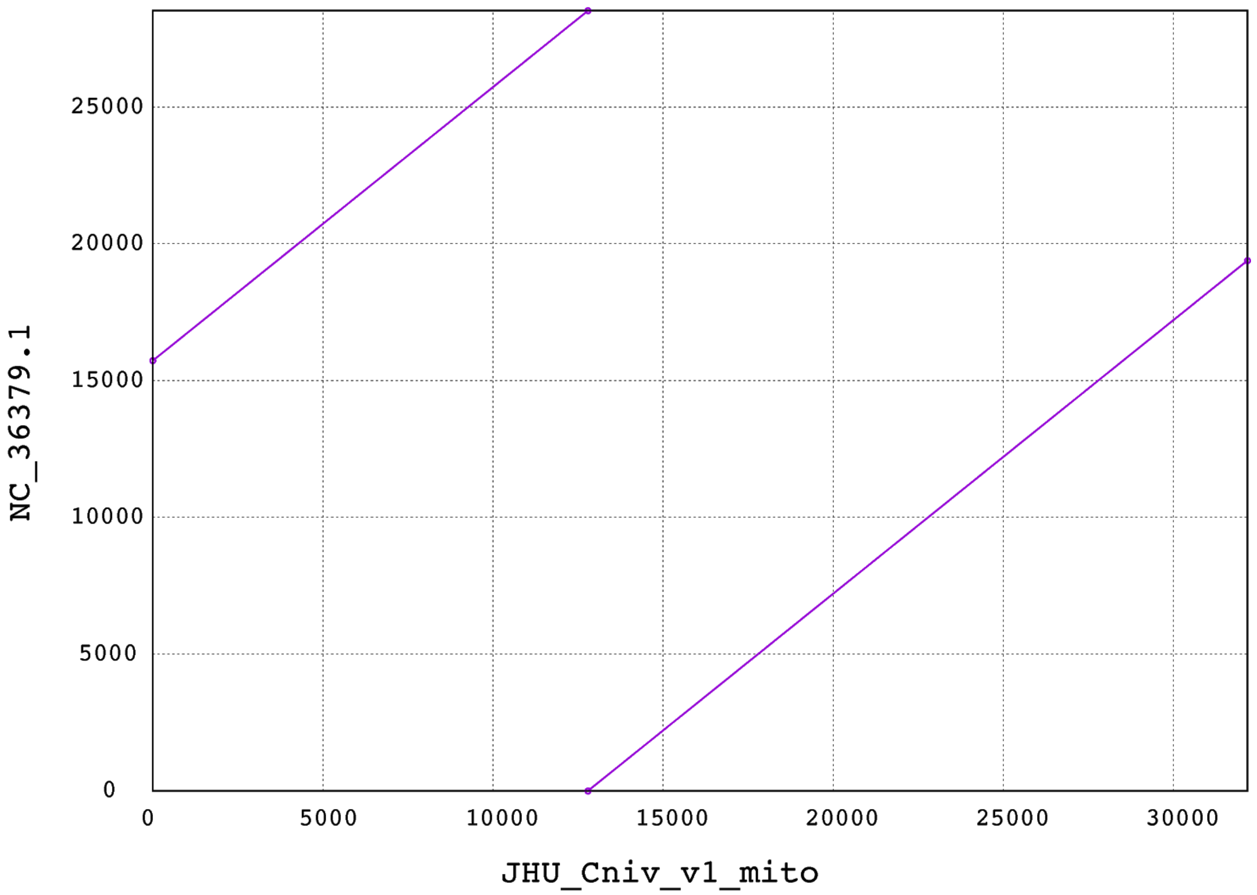


**Supplemental Figure 2**: Alignment of our 32Kb circular contig (x axis) with the completed mitochondrial genome of the *C. nivariensis* reference genome (y axis). The final 3662bp of this contig appears twice in the reference genome.

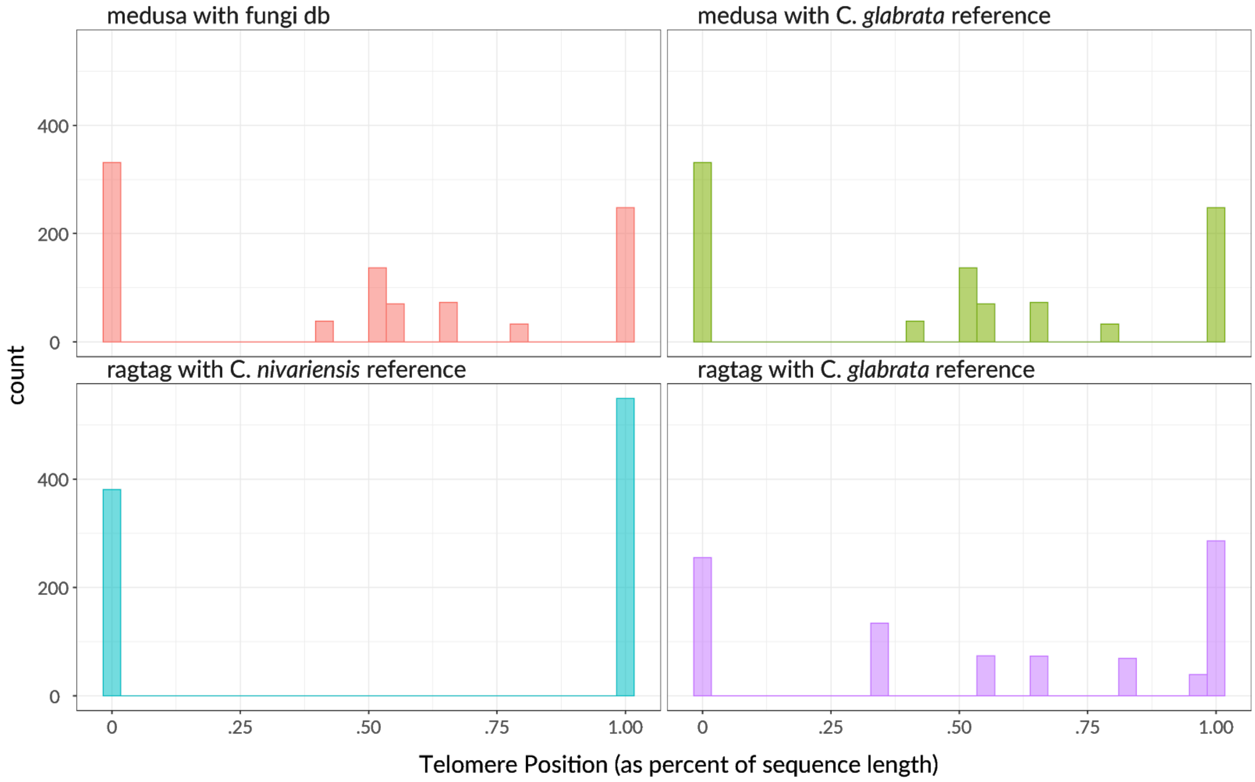


**Supplemental Figure 3:** Histogram of telomere repeat positions in our assembly, and in scaffolds produced by RagTag and MeDuSa. When MeDuSa is used with a database including the reference genomes of C. *nivariensis*, C. *glabrata*, C. *bracarensis*, and N. *delphensis*, telomeres are placed in the middle of contigs. The same result is produced when only the C. *glabrata* genome is used for scaffolding with MeDuSa, and MeDuSa fails to run when only the C. *nivariensis* reference is used. When the *C. nivariensis* reference genome is used for scaffolding with RagTag, no changes are made. When the more contiguous *C. glabrata* genome is used with RagTag, telomere sequences are again placed in the middle of sequences, suggesting a scaffolding error.

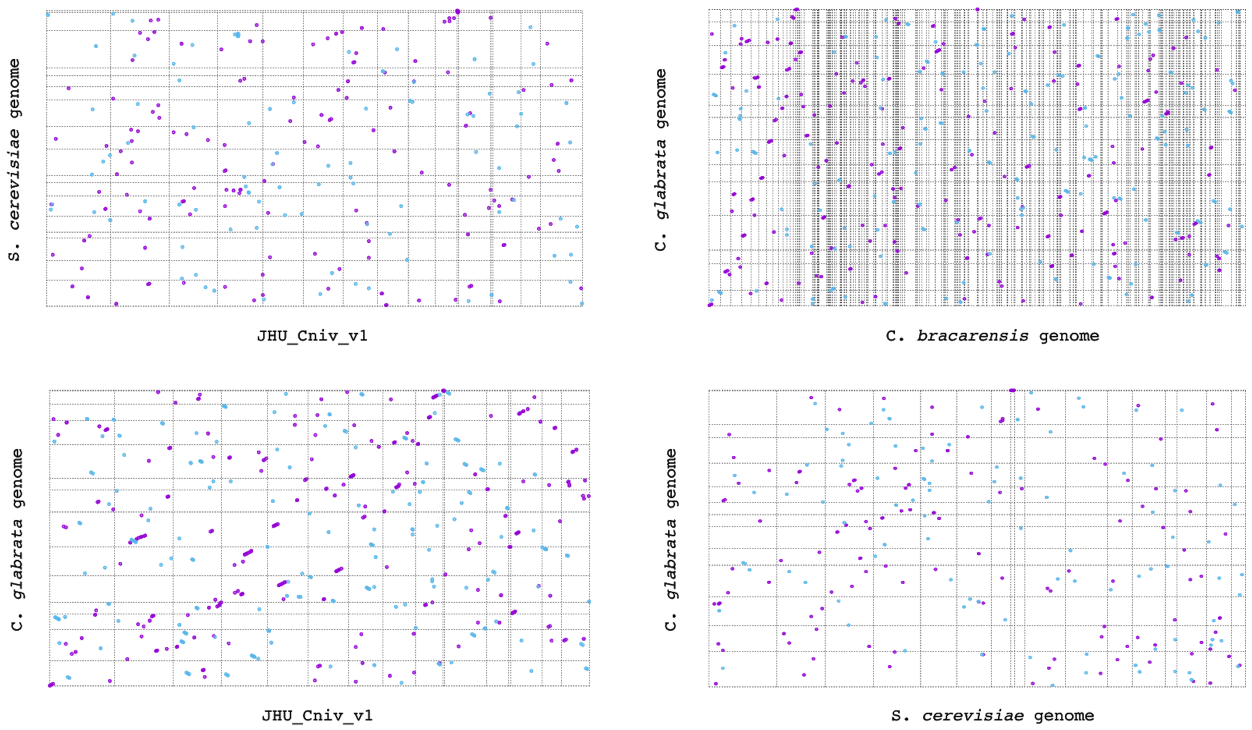


**Supplemental Figure 4:** Whole genome alignment of our new assembly against the S. *cerevisiae* (top left), and C*. glabrata* (bottom left) reference genomes. For both, there are no long alignments, suggesting that there is little similarity in genome structure between these species and *C. nivariensis*. C. *bracarensis*, a close relative to both C. *glabrata* and C. *nivariensis*, also shares little genome similarity to C. *glabrata* (top right), suggesting that yeast genomes within the glabrata clade are not generally similar enough to support inter-species reference based scaffolding. We also compared C. *glabrata* to the highly contiguous and complete S. *cerevisiae* genome (bottom right) to check that genome contiguity alone did not bias the genome similarity detected.

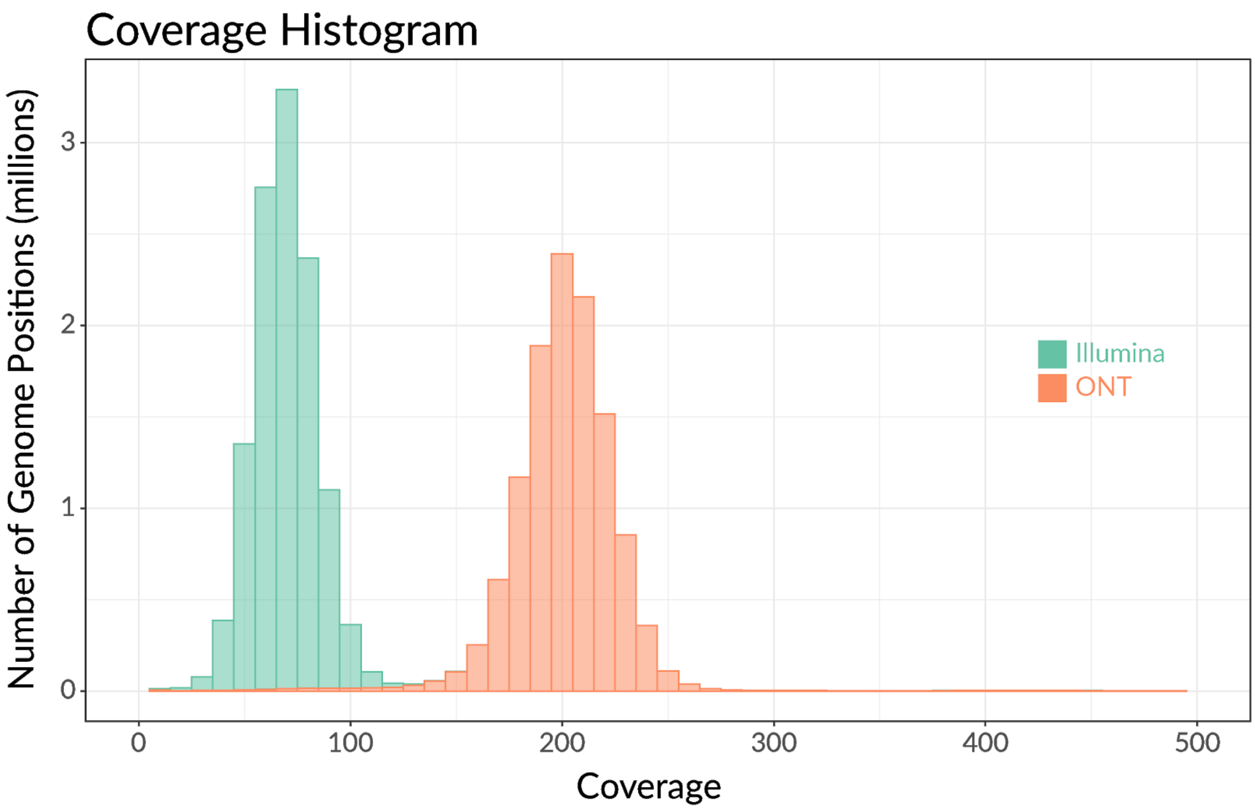


**Supplemental Figure 5**: Histogram of coverage per base in our assembly by filtered (>3kb) ONT reads and trimmed Illumina reads.

| Contig | Length (bp) | Forward Telomeres | Reverse Telomeres |
|---|---|---|---|
| tig01 | 1423475 | 35 | 38 |
| tig02 | 1283968 | 0 | 39 |
| tig03 | 1060011 | 35 | 39 |
| tig04 | 933062 | 36 | 26 |
| tig05 | 1010854 | 0 | 36 |
| tig06 | 885783 | 35 | 38 |
| tig07 | 879540 | 39 | 35 |
| tig08 | 763992 | 34 | 33 |
| tig09 | 714796 | 35 | 47 |
| tig10 | 675194 | 36 | 36 |
| tig11 | 594828 | 32 | 26 |
| tig12 | 617546 | 36 | 0 |
| tig13 | 481613 | 38 | 41 |
| tig14 | 434809 | 33 | 33 |
| tig24 | 44616 | 0 | 39 |
| JHU_Cniv_v1_mito | 28512 | 0 | 0 |

**Supplemental Table 1:** Contig lengths and the number of times the forward and reverse telomere sequence appears in each.

| | Total | Gene | Exon |
|---|---|---|---|
| Augustus (BRAKER) | 23,497 | 5,028 | 6,109 |
| Genemark.hmm (BRAKER) | 36 | 6 | 12 |
| Liftoff glabrata | 263 | 130 | 2 |
| Liftoff cerevisiae | 42 | 21 | 0 |
| Liftoff albicans | 0 | 0 | 0 |
| StringTie | 2,141 | 824 | 1,175 |

**Supplemental Table 2:** Annotation contributions from each software

| | Total Exons | Total Genes |
|---|---|---|
| JHU_Cniv_v1 | 7,298 | 5,859 |
| C. *glabrata* | 5,629 | 5,448 |
| S. *cerevisiae* | 6,760 | 6,420 |
| C. *albicans* | 6,732 | 6,263 |

**Supplemental Table 3:** Gene and exon counts of our annotation and currently available reference annotations.
